# Sphingolipids mediate polar sorting of PIN2 through phosphoinositide consumption at the *trans*-Golgi Network

**DOI:** 10.1101/2020.05.12.090399

**Authors:** Yoko Ito, Nicolas Esnay, Matthieu Pierre Platre, Lise C. Noack, Wilhelm Menzel, Stephane Claverol, Patrick Moreau, Yvon Jaillais, Yohann Boutté

**Affiliations:** Laboratoire de Biogenèse Membranaire, UMR5200, Université de Bordeaux, CNRS, 33140 Villenave d’Ornon, France; Laboratoire Reproduction et Développement des Plantes, Université de Lyon, ENS de Lyon, UCB Lyon1, CNRS, INRAE, F-69342 Lyon, France; Proteome Platform, Functional Genomic Center of Bordeaux, Université de Bordeaux, F-33076, Bordeaux, France; Bordeaux Imaging Center, UMS 3420, Université de Bordeaux, CNRS, 33000 Bordeaux, France

**Author notes:** These authors equally contributed to the work (Yoko Ito, Nicolas Esnay). BioDiscovery Institute and Department of Biological Sciences, University of North Texas, Denton, TX 76203, USA. Plant Molecular and Cellular Biology Laboratory and Integrative Biology Laboratory, Salk Institute for Biological Studies, La Jolla, United States (Matthieu Pierre Platre).

## Abstract

The lipid composition of organelles acts as a landmark to define membrane identity and specify subcellular function. Phosphoinositides are anionic lipids acting in protein sorting and trafficking at the *trans*-Golgi network (TGN). In animal cells, sphingolipids are known to control the turnover of phosphoinositides through lipid exchange mechanisms at endoplasmic reticulum/TGN contact sites. In this study, we discovered a completely new mechanism acting on sphingolipid-mediated phosphoinositides homeostasis at the TGN in plant cells. We used multi-approaches to show that a reduction of the acyl-chain length of sphingolipid results in increased level of phosphatidylinositol-4-phosphate (PI4P) at the TGN, independently from either lipid exchange induced by sphingolipid synthetic flux, or local PI4P synthesis. Instead, we found that sphingolipids mediate the consumption of PI4P through phosphoinositide-specific phospholipase C (PI-PLC) and this process impacts the sorting of the auxin efflux carrier PIN2 at the TGN. Together, our data identify a new mode of action of sphingolipids in lipid interplay at the TGN during protein sorting.

---

Post-Golgi protein sorting is a fundamental process to direct proteins to polar domains of eukaryotic cells^1–4^. The *trans*-Golgi Network (TGN) is an essential organelle acting in cargo sorting. TGN malfunctioning results in serious diseases as well as cell polarity, differentiation and organ development defects in both animal and plant kingdoms^3–5^. Lipid interplay between sphingolipids (SL), sterols and phosphoinositides (PIPs), is thought to orchestrate sorting and trafficking of secretory cargos at the TGN^6–9^. SL and sterols are enriched at the TGN where they increase membrane thickness and assemble to form small sorting platforms, both of which are important for protein sorting and trafficking^10–14^. Importantly, SL metabolic flux controls PIP homeostasis through PIPs/sterols or PIPs/PS exchanges between the ER and the TGN^8,9,15^. This effect of SL over PIPs is crucial as PIPs favor vesicle budding and fission and act in polarized trafficking in concert with small GTPases or elements of the exocyst complex^7,16–19^. Additionally, PIPs recruit adaptor proteins or membrane curvature-sensitive proteins that help selecting cargos and forming vesicles^20–23^. In animal cells, the transfer of phosphocholine from phosphatidylcholine (PC) to ceramide occurs at the TGN and produces sphingomyelin and a diacylglycerol (DAG) molecules that favor negative membrane curvature and fission, and activates the PI4KinaseIIIβ (PI4KIIIβ) which locally produces PI-4-phosphate (PI4P)^8,21,24,25^. PI4P recruits both CERT, facilitating ceramide transfer, and oxysterol-binding proteins (OSBP), which exchange PI4P for sterols at ER-TGN contact sites^9,26^. This process negatively feedbacks on OSBP localization at the TGN and the transfer of ceramide from the ER to the TGN. Hence, there is a homeostatic control of SL synthetic flow over PI4P turnover at ER/TGN contact sites which is dependent on PI4P consumption through OSBP-mediated sterol exchange and proximity of OSBP with PI4K^8,9^. Some OSBP-related proteins (ORPs) operate a phosphatidylserine (PS)/PI4P exchange at the TGN rather than sterol/PI4P exchange^15^. In plants, OSBP proteins have been evidenced at the ER to *cis*-Golgi interface^27^. However, no ER/TGN membrane contact sites have been unambiguously shown so far in plants, probably due to the highly dynamic nature of the TGN which can detach from Golgi apparatus and become an independent organelle^28,29^. Nonetheless, SL and sterols are enriched at the TGN^30,31^. Moreover, PI4P and PS participate to establish an electrostatic territory at the plant TGN^32,33^. Interestingly, alteration of either the acyl-chain length of SL or PI4KIIIβ function result in swollen TGN-vesicles being less interconnected with membrane tubules and alteration in sorting of the auxin efflux carrier PIN2 at the TGN, indicating a potential interplay between SL and PIPs at the TGN in plant cells^30,34^. We employed a combination of live cell biology, immuno-purification of targeted TGN compartments, label-free proteomics and lipid biochemistry approaches to now reveal that the acyl chain length of SL plays a role in PIPs homeostasis and sorting of PIN2 at the TGN, independently from sterols or PS homeostasis or from PIP-related kinases or phosphatases. Unexpectedly, we identified that the acyl-chain length of SL impacts on PI4P consumption at the TGN through PIPs-specific phospholipase C (PI-PLC) and this lipid interplay plays a role in sorting of the auxin efflux carrier PIN2 at the TGN. Altogether, our results establish a completely new mode of action of SL on PIPs homeostasis during protein sorting through another mode of PI4P consumption than the synthesis-induced PI4P/sterols exchange mechanism.

## Results

### PI4P, but not PS, quantity at TGN, is modulated by the acyl-chain length of SL

In *Arabidopsis* root epidermal cells, PI4P mainly resides at the plasma membrane (PM) and sparsely at the TGN/EEs^32,35^. To analyze the potential effect of the acyl-chain length of SL on the localization of PI4P, we used metazachlor, a chemical allowing a fine-tunable reduction of C24- and C26-α-hydroxylated fatty acids (FAs), h24 and h26, which are specifically present in the pool of SL, without modifying the total quantity of SL^30^. We now show that shortening the acyl chain length of SL by metazachlor treatment increases PI4P quantity in the membrane of intracellular compartments (Fig. 1a, b), as revealed by the *in vivo* genetically encoded biosensor 1x PH domain of the Human FAPP1 protein fused to mCITRINE (mCIT-1xPH^FAPP1^)^35^. As compared to untreated roots, 50 nM or 100 nM metazachlor treatment resulted in increased signal intensity in intracellular dots (Fig. 1b). The PH domain of the FAPP1 protein binds to both PI4P and the small GTPase ARF1 which partly localizes at the TGN^36,37^. Hence, we checked that the increase of signal observed upon metazachlor was not due to increased binding of the mCIT-1xPH^FAPP1^ sensor to the ARF1 protein. We used the mCIT-1xPH^FAPP1-E50A^ and mCIT-1xPH^FAPP1_E50A_H54A^ sensors mutated for ARF1 binding but not PI4P binding^32^. Consistent with previous observations in *Nicotiana benthamiana*^32^, we could hardly detect any signal at the TGN in both mCIT-1xPH^FAPP1-E50A^ and mCIT-1xPH^FAPP1-E50A-H54A^ untreated roots cells (Fig. 1d and Supplementary Fig. 1a). In contrast, the signal of both biosensors was clearly increased in intracellular compartments in metazachlor treated cells (Fig. 1d, e and Supplementary Fig. 1a, b). In addition, the intensity of ARF1-GFP in dots was not altered upon metazachlor (Supplementary Fig. 1d, e). Hence, we excluded the possibility that the increase of fluorescence intensity of the 1xPH^FAPP1^ PI4P sensor was due to higher binding to ARF1 or more ARF1 at the TGN upon metazachlor. To reinforce our data, we analyzed PI4P sensor lines with higher number of PH domains, *i.e*. mCIT-2xPH^FAPP1^ and mCIT-3xPH^FAPP1^ which increase the avidity of the sensor to PI4P^32^. Consistent with previously published results, both mCIT-2xPH^FAPP1^ and mCIT-3xPH^FAPP1^ sensors display strong PM labeling and almost undetectable levels of fluorescence at the TGN^32,35^ (Fig. 1g, j). Strikingly, both mCIT-2xPH^FAPP1^ and mCIT-3xPH^FAPP1^ showed clear increase of signal intensity in intracellular compartments upon metazachlor (Fig. 1h, k). We also checked the fluorescence intensity of PI4P sensors at the PM and observed that mCIT-1xPH^FAPP1^ intensity at the PM was decreased upon metazachlor (Fig. 1c). Contrastingly, mCIT-2xPH^FAPP1^ and mCIT-3xPH^FAPP1^ intensity were not decreased at the PM upon metazachlor (Fig. 1i, l). These results are consistent with the previous observation that mCIT-2xPH^FAPP1^ and mCIT-3xPH^FAPP1^ dwell time is higher than mCIT-1xPH^FAPP1^ in PI4P-riched membranes due to higher avidity of the sensors^32^. Similarly, mCIT-1xPH^FAPP1-E50A^ and mCIT-1xPH^FAPP1-E50A-H54A^ intensity at the PM were not decreased upon metazachlor (Fig. 1f and Supplementary Fig. 1c). To confirm our observation in mCIT-1xPH^FAPP1^ sensor line, we analyzed another PI4P biosensor that strictly labels the pool of PI4P at the PM *i.e*. the P4M domain of the *Legionella pneumophila* SidM protein fused to mCITRINE^32^ (mCIT-P4M^SidM^). Our results were similar to mCIT-1xPH^FAPP1^ sensor, the intensity at the PM was decreased upon metazachlor treatment (Supplementary Fig. 1f, g). Hence, the effect of metazachlor on the pool of PI4P at the PM appears to be slighter as compared to the consistent increase we observed at the TGN. It was previously shown that PS acts in concert with PI4P to generate an electrostatic gradient between the PM and the TGN^33^. Hence, we wondered whether the increase of PI4P upon metazachlor would also be visible for PS. We used the C2 domain of bovine Lactadherin fused to mCITRINE (mCIT-C2^LACT^) to visualize PS^33^. Our results did not evidence any change in PS fluorescence intensity at the PM or in intracellular compartments upon metazachlor treatment (Fig. 1m-o). This indicates that the acyl chain length of SL has some specificity and does not target all anionic lipids. Moreover, our results further indicate that the increase of PI4P at the TGN upon metazachlor is unlikely to be correlated with a PS/PI4P exchange mechanism.

**Fig. 1.**
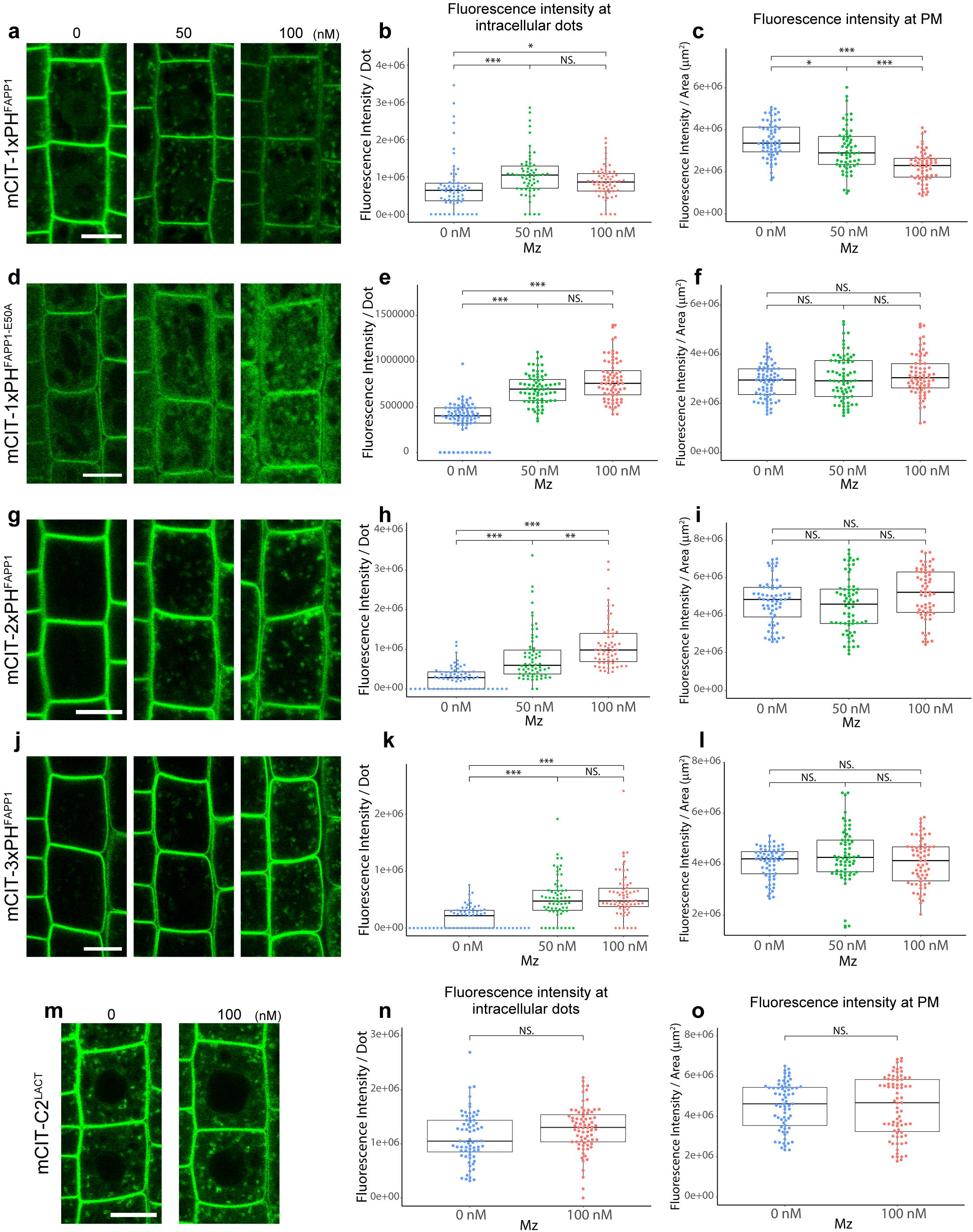
The acyl-chain length of SL is required for the intracellular distribution of PI4P, but not PS. Confocal micrographs of *Arabidopsis* root epidermal cells expressing either the PI4P biosensor mCIT-1xPH^FAPP1^ (a), mCIT-1xPH^FAPP1-E50A^ (d), mCIT-2xPH^FAPP1^ (g), mCIT-3xPH^FAPP1^ (j), or the PS biosensor mCIT-C2^LACT^ (m) upon 0, 50, or 100 nM metazachlor (Mz) treatment. Fluorescent intensity at intracellular dots (b, e, h, k, n) and at PM (c, f, i, l, o) was quantified (n = more than 60 cells distributed over at least 20 roots for each condition). All the PI4P biosensors tested showed significant increase of the signal intensity at intracellular dots upon Mz treatment, and some of them showed a decrease of the signal at PM (a-l). In contrast, the distribution of the PS biosensor was not affected (m-o). Statistics were done by two-sided Wilcoxson’s rank-sum test, \**P*-value<0.01, ** *P*-value <0.001. *** *P*-value <0.0001. Scale bars, 10 μm.

### The acyl-chain length of SL acts on PI4P accumulation at the TGN independently of PI4P synthesis or a PI4P-sterol exchange mechanism

Increased PI4P level at intracellular compartments could be due to more synthesis through the PI4Kinases (PI4K) activity. To test this hypothesis, we treated *Arabidopsis* seedlings with wortmannin (Wm) that inhibits both PI4K and PI3K at 33 μM^38–40^. In the control seedlings, PI4P, labelled by the mCIT-3xPH^FAPP1^ biosensor, was still mainly localized at PM and weakly in intracellular dots after 5 min of Wm treatment (5 min) (Fig. 2a). However, PI4P intensity at the PM already decreased after 15 min of Wm treatment confirming the efficiency of Wm treatment (Fig. 2b). In metazachlor grown seedlings, PI4P was accumulated in dots at the beginning of the Wm treatment (5 min and 15 min), as in metazachlor treated seedlings without Wm (Fig. 2c). Notably, this accumulation of PI4P sensor in dots decreased only slightly and was statistically insignificant after 90 min of Wm treatment (Fig. 2c), indicating that PI4K is not primarily involved. Our results indicate that metazachlor effect on PI4P is not dependent on PI4K activity.

**Fig. 2.**
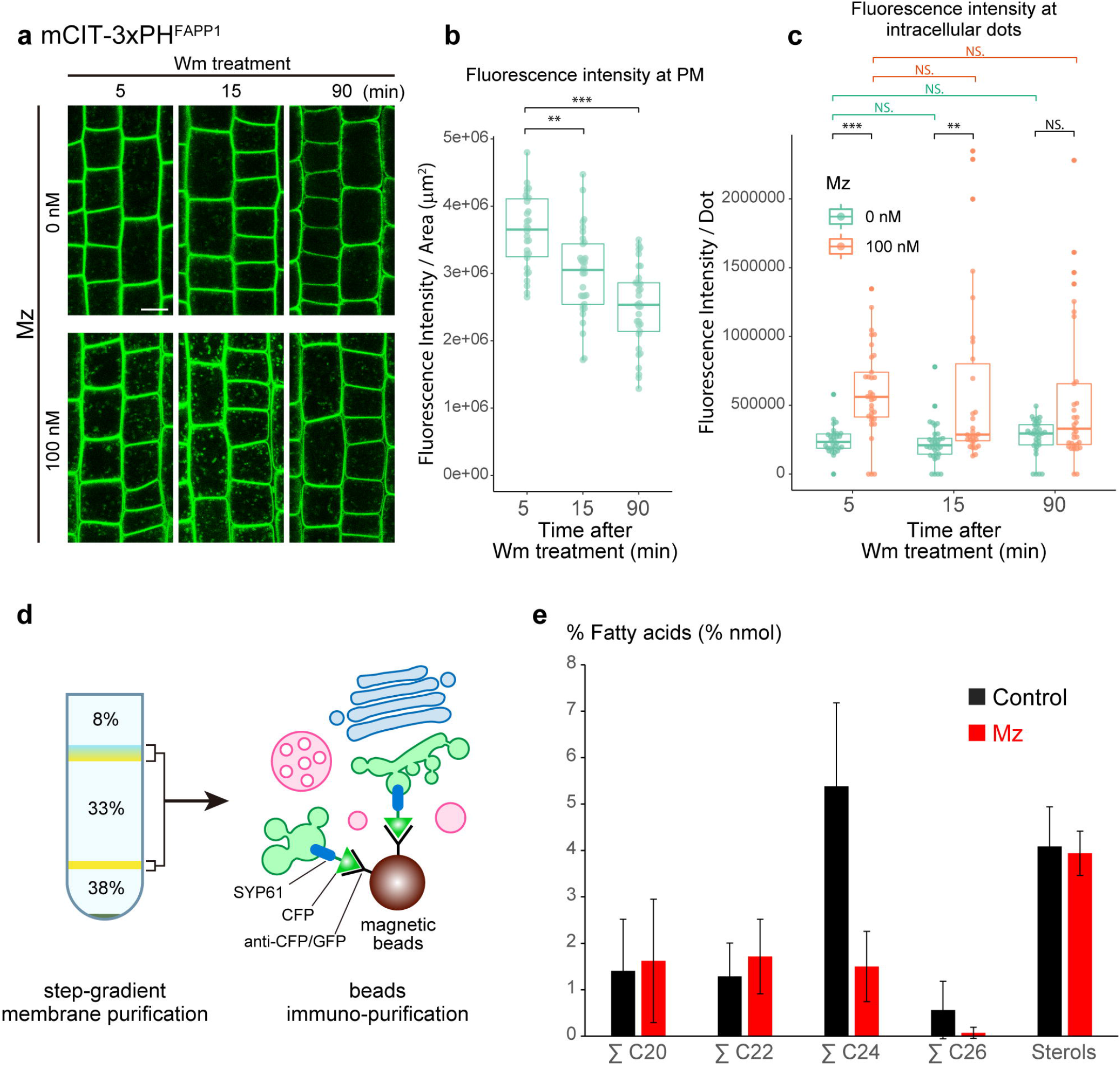
Neither PI4P synthesis by PI4K nor PI4P-sterol exchange is involved in SL-mediated PIP homeostasis. (a) Confocal images of root epidermal cells expressing the PI4P biosensor mCIT-3xPH^FAPP1^ during Wm treatment (5, 15 and 90 min) of seedlings grown on 0 or 100 nM Mz-containing medium. (b) Quantification of fluorescence intensity at the PM of the cells upon Wm treatment alone (n = more than 30 cells distributed over at least 10 roots for each condition). The PI4P pool at the PM was decreased significantly. (c) Quantification of the fluorescence intensity at intracellular dots (n = more than 30 cells distributed over at least 10 roots for each condition). The PI4P pool accumulated at intracellular dots by Mz was not significantly decreased upon Wm treatment. (d) Immuno-purification of intact SVs/TGN membrane compartments. Step-gradient-purified membrane fraction was incubated with magnetic beads conjugated with CFP/GFP antibodies, and the intact compartments labeled by the SVs/TGN marker SYP61-CFP were immuno-purified. (e) GC-MS analysis of fatty acid and sterol composition at immunopurified SYP61-SVs/TGN fraction in control condition (black) or treated with 100 nM Mz (red) (n = 3 biological replicates, error bars are s.d.). While Mz strongly decreased the sum of C24 and the sum of C26 fatty acids it did not affect the sterol content at SVs/TGN. Statistics were done by two-sided Wilcoxson’s rank-sum test, ** *P*-value <0.001, *** *P*-value <0.0001. Scale bars, 10 μm.

In animal cells, SL flow regulates PI4P homeostasis at the TGN through an OSBP-mediated PI4P/sterol exchange mechanism^8,9,21,23,24,26^. We reasoned that if this would be the case in plant cells, an SL-mediated accumulation of PI4P at the TGN would imply an equivalent decrease of sterols at the TGN. We immunopurified SYP61-SVs/TGN compartments (Fig. 2d) and characterized the fatty acid (FA) and sterol composition by GC-MS in untreated and metazachlor-treated seedlings. Our results revealed that, in SVs/TGN IPs, metazachlor strongly decreases C24- and C26-FAs (Fig. 2e), mainly in the pool of α-hydroxylated FAs (hFAs) which are specific to SLs (Supplementary Fig. 2a), consistently to what we published before on whole *Arabidopsis* roots lipid composition^30^. Moreover, metazachlor did not alter the C16- and C18-FAs which are a hallmark for phospholipids (Supplementary Fig. 2b and c). Contrastingly to h24 and h26 FAs, the sterol content in SYP61-SVs/TGN IP compartments was not significantly modified upon metazachlor (Fig. 2e). These results argue for a role of SL on PI4P homeostasis independent from a sterol exchange mechanism.

### The abundance of PIP-specific phospholipase C at SYP61-SVs/TGN is modulated by the acyl-chain length of SL

To identify proteins whose abundance at the TGN is sensitive to the acyl-chain length of SL, we performed label-free quantitative proteomics by LC-MS/MS on four biological replicates of SYP61-SVs/TGN IP compartments (Fig. 2d) from control and metazachlor-treated seedlings. To check the efficiency of immuno-purification we loaded equal amount of IP input and output fractions on SDS-PAGE (Supplementary Fig. 3a) and performed western-blotting with an anti-GFP antibody. Our results showed that the IP output fraction was enriched for SYP61-CFP as compared to the IP input fraction (Supplementary Fig. 3b). Moreover, the SVs/TGN-resident protein ECHIDNA was also found to be enriched in the IP output fraction (Supplementary Fig. 3c). These results confirmed the efficient purification of SYP61-SVs/TGN compartments. We only kept proteins that were consistently identified in each of the four biological replicates, resulting in a list of 4428 proteins. Due to the genetic redundancy in some protein families, some peptides could correspond to either one accession or the sum of several ones (indicated in Supplementary Data 1). Although the label-free methodology aims at comparing protein abundancies of a sample upon different conditions, general observations can be drawn from individual protein abundancies. Indeed, based on mass spectrometry abundance measurement, SYP61 was one of the most abundant protein, confirming the efficiency of the IP (Supplementary Data 1). We checked the abundance of known SVs/TGN proteins and found high abundance of the V-ATPase VHA-a1^41^, the RAB-GTPase-interacting YIP4a/b proteins and its interacting partner ECHIDNA^42^, small GTPases of the ARF1 family, the vacuolar protein sorting45 VPS45 and its interacting proteins SYP43, SYP42 and VTI12^43–45^ (Fig. 3a, Supplementary Data 2). SYP41 was found at much lower abundance than SYP43 or SYP42 indicating that SYP41 localizes at a distinct sub-domain of TGN than SVs/TGN, consistently with previous work^43,45^ (Fig. 3a, Supplementary Data 2). Small GTPases of the RAB family, RAB-A4b and RAB-A5d, described to localize at SVs^46^, were present (Fig. 3a, Supplementary Data 2). We could not find Golgi proteins MEMBRIN11/12, SYP32 and GOT1, confirming the purity of SVs/TGN fraction over the Golgi (Fig. 3a, Supplementary Data 1). We checked the MVB/LE markers RAB-F2b, RAB-F2a and RAB-F1 and found them in low abundance with RAB-F2b being the most present consistently to its dual MVB and TGN localization described previously^47^ (Fig. 3a, Supplementary Data 2). Interestingly, none of these proteins were significantly down- or up-regulated at SVs/TGN upon metazachlor (Fig. 3b, Supplementary Data 2). Next we checked SL-related enzymes and found low abundance of the SL desaturase SLD1, intermediate abundance of the SL kinases SPHK1/2 and LCKB2, the alkaline and neutral ceramidases ACER and NCER1/2/3, the inositolphosphorylceramide (IPC) Synthase2 (IPCS2) that grafts inositolphosphate group on ceramide to produce IPC^48^, high abundance of the inositolphosphorylceramide glucuronosyltransferase1 (IPUT1) enzyme that grafts a glucuronic acid (GlcA) on IPC^49^ to produce GlcA-glucosylinositolphosphorylceramides (GlcA-GIPCs), and intermediate abundance of the GIPC mannosyltransferase1 (GMT1) enzyme that grafts mannose on GlcA-GIPC^50^ to produce final GIPC (Supplementary Fig. 3d, Supplementary Data 2). Since the main form of SL are GIPC in plants, our results suggest that SVs subdomain of TGN is a main place for SL synthesis, consistently with previous localization of GIPC enzymes at the Golgi complex and enrichment of SL at SVs/TGN^30,48–50^. However, none of these SL-related proteins were modified at SVs upon metazachlor (Supplementary Fig. 3f, Supplementary Data 2). We did not detect any OSBP proteins or OSBP-related proteins (ORP) in agreement with a metazachlor effect independent from a sterol-exchange mechanism. We further checked PS-related enzymes and found low abundance of the PS synthase1 (PSS1), intermediate abundance of two PS flippases (ALA1 and ALA2), one PS decarboxylase (PSD3) and high abundance of two PS/PE-specific phospholipase D (PLD-gamma1/2/3 and PLD-beta1/2) (Supplementary Fig. 3e, Supplementary Data 2). None of these PS-related proteins were modified at SVs upon metazachlor treatment consistently to the unaltered localization of PS biosensors (Supplementary Fig. 3f, Supplementary Data 2). Finally, we checked the abundance of PIP-related proteins and found low abundance of the PI synthases PIS1/2, the PI4Kinase PI4α1, the PI(3,5)P2 phosphatases SAC1-5, the PI4P phosphatase SAC8, the PI(4,5)P2 phosphatases IP5P1 and SAC9, the PI3P and PI(3,5)P2 phosphatases PTEN2A and PTEN2B, and high abundance of the PI4P phosphatases SAC6/7 (Fig. 3c, Supplementary Data 2). SAC6 is specific to flowers, as we performed the proteomics on seedlings we believe that SAC7 is the most abundant at SVs/TGN which is consistent with previous observation that SAC7/RHD4 localizes at post-Golgi structures in *Arabidopsis* root hair cells^51^. Importantly, none of these proteins were modified upon metazachlor treatment indicating that SLs do not target localization of PIP-related phosphatases, more particularly PI4P phosphatases SAC6/7 that are the most present at SVs/TGN (Fig. 3e, Supplementary Data 2). Interestingly, we found proteins of the PIPs-specific phospholipase C family (PI-PLC) that hydrolyze PIPs to produce DAG and inositol polyphosphate^52^. We found two low abundant PI-PLCs-X domain-containing proteins and high abundance of PI-PLC2/7 proteins (Fig. 3d). Both PI-PLCs-X and PI-PLC2/7 were strongly reduced at SVs/TGN upon metazachlor (Fig. 3e). It was shown previously that PLCs hydrolyze equally well both PI4P and PI(4,5)P2 *in vitro* and that genetic depletion of *PLCs* results in higher amount of PI4P and PI(4,5)P2, suggesting that PI4P and PI(4,5)P2 are the *in planta* substrates of PI-PLCs^53–55^. Hence, a lower amount of PI-PLCs at SVs/TGN could result in a higher amount of PI4P at the TGN.

**Fig. 3.**
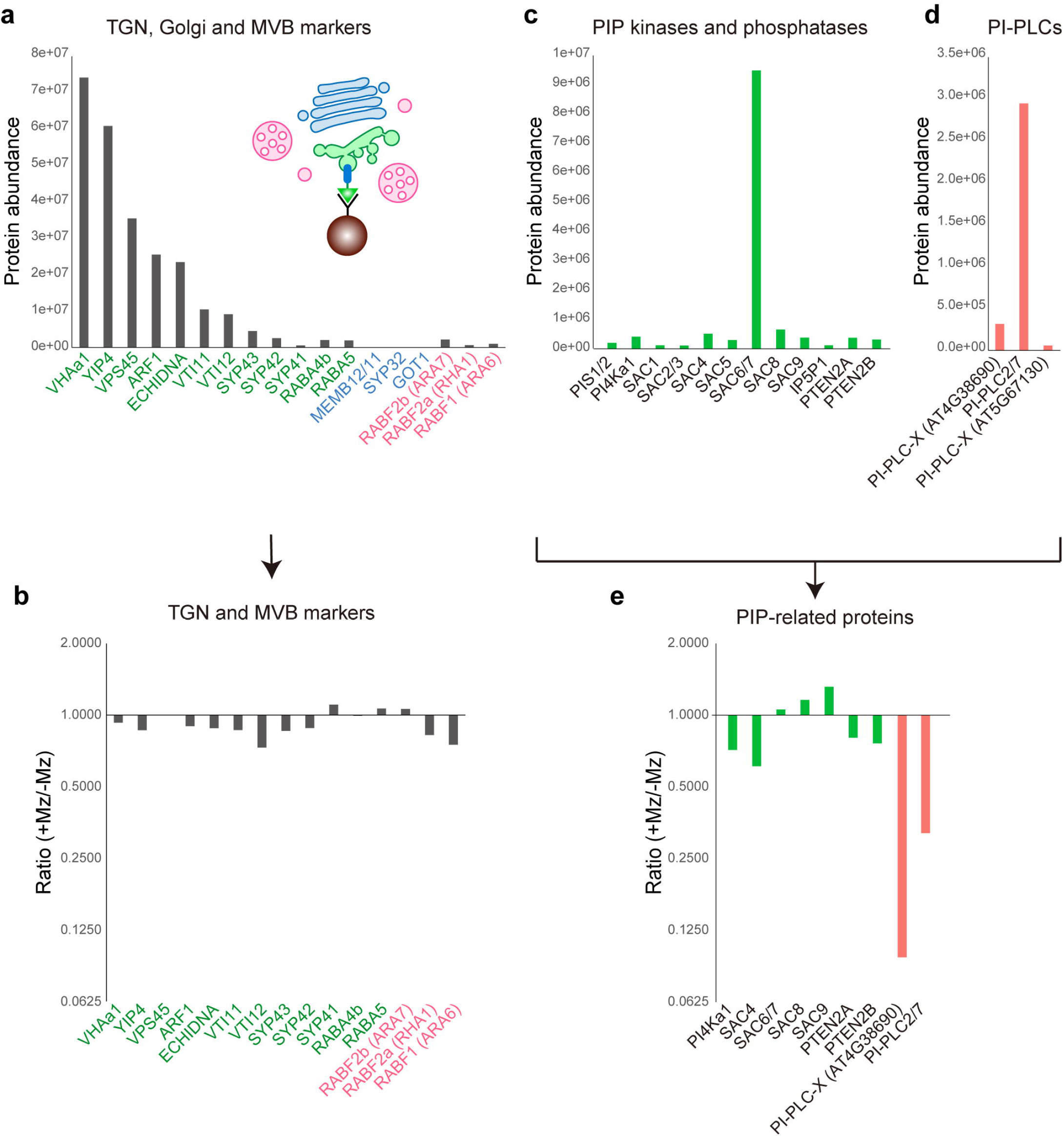
LC-MS/MS label-free proteomics identification of proteins which abundance at SVs/TGN is sensitive to acyl-chain length of SL. (a, c, d, e) Protein abundance in SYP61 SVs/TGN immuno-purified compartments analyzed by label-free quantitative proteomics (detailed analysis is synthesized in Supplementary Data 2, including the number of peptides found for each proteins). (a) TGN (green), Golgi (blue) and MVB (red) markers, (c) PIP kinases and phosphatases, (d) PI-PLCs. The abundance ratio of the proteins shown in (a) or (c, d) between control and Mz treated samples is displayed in (b) or (e), respectively. For the calculation of ratio, we applied a threshold of minimal protein amount as 300,000 to select the proteins from which we can get reliable ratio. TGN markers were abundant compared to Golgi and MVB markers, which indicates an efficient purification, and their abundance was not affected by Mz treatment. In contrast, PI-PLCs were strongly reduced upon Mz treatment.

### SL-mediated PI4P homeostasis involves PI-PLCs and impacts PIN2 sorting at SVs/TGN

To explore a potential implication of PI-PLCs in metazachlor-induced accumulation of PI4P at the TGN, we used the mCIT-2xPH^FAPP1^ PI4P biosensor upon 90 min treatment with either the PI-PLC inhibitor U73122 or its inactive analog U73343 at 1 μM and 5 μM for both active and inactive analogs^56–58^ (Fig. 4a). We did not observe any significant change in fluorescence intensity at intracellular dots between 1 μM and 5 μM of inactive U73343 control treatment (Fig. 4b). Contrastingly, the active U73122 PI-PLCs inhibitor treatment clearly displayed a significant increase of PI4P at 1 μM and this effect was further increased at 5 μM (Fig. 4b). These results revealed that PI-PLCs play a role on PI4P homeostasis in intracellular compartments. Furthermore, when seedlings were grown on 100 nM metazachlor prior to treatment with 1 μM of active U73122, the pool of PI4P in intracellular compartments got even stronger (Fig. 4b). However, with 5 μM of active U73122 PI-PLCs inhibitor, there was no significant difference between seedlings grown on either 0 or 100 nM metazachlor (Fig. 4b). These results argue that the acyl-chain length of SLs acts in PI4P accumulation at the TGN through the modulation of a pool of PI-PLC proteins. It was shown previously that PI4P mainly accumulates at the PM with a minor pool at the TGN^32,33^. Hence, it is expected that the PI4P accumulation upon SLs modification by metazachlor or PI-PLC inhibition by U73122 occurs at the TGN. To unambiguously identify the nature of the intracellular compartments in which PI4P accumulates, we performed co-localization of the mCIT-2xPH^FAPP1^ PI4P biosensor with the SVs/TGN marker VHA-a1 fused to mRFP^59^. Our results revealed a strong colocalization between 2xPH^FAPP1^ and VHA-a1 upon metazachlor treatment, confirming that the intracellular accumulation of PI4P occurs at SVs/TGN when the acyl chain length of SLs is reduced (Fig. 5a, e). Although weaker, we identified a good level of co-localization between 2xPH^FAPP1^ and VHA-a1 upon inhibition of PI-PLC by U73122 treatment as well (Fig. 5a, e). Both PI-PLCs and acyl-chain length of SLs are known to be involved in root gravitropism, auxin distribution and PIN2 apical polarity^30,56,60,61^. However, unlike acyl-chain length of SLs, whether PI-PLCs are involved in PIN2 sorting at SVs/TGN is not known. Hence, we thoroughly quantified the fluorescence intensity of PIN2, specifically at intracellular dots, upon inhibition of PI-PLC by active U73122 treatment or upon treatment with its inactive analog U73343. Our results revealed no difference between the control condition and treatment with the inactive PI-PLC inhibitor analog U73343 (Fig. 5b, c). However, a strong accumulation of PIN2 in intracellular compartments was observed upon treatment with 5 μM of active PI-PLC inhibitor U73122 (Fig. 5b, c). To identify the compartments in which PIN2 accumulates, we performed co-localization experiments between PIN2 and the SVs/TGN-localized SYP61 syntaxin and VTI12 SNARE markers. The level of co-localization we quantified between PIN2 and SVs/TGN markers was similar to what we quantified between 2xPH^FAPP1^/PI4P and the SVs/TGN marker VHA-a1 upon the active PI-PLC inhibitor U73122 (Fig. 5d, e). Importantly, we quantified a weak co-localization between PIN2 and the Golgi-localized SYP32 syntaxin and MEMBRIN12 SNARE markers upon U73122 (Fig. 5d, e). Altogether, our results consistently support a role of PI-PLCs in mediating the effect of SLs on the accumulation of PI4P at SVs/TGN, concomitantly with an accumulation of the auxin efflux carrier PIN2 at SVs/TGN. Hence, our results suggest that SLs act on PIN2 sorting at SVs/TGN through the modulation of PI4P consumption by PI-PLCs.

**Fig. 4.**
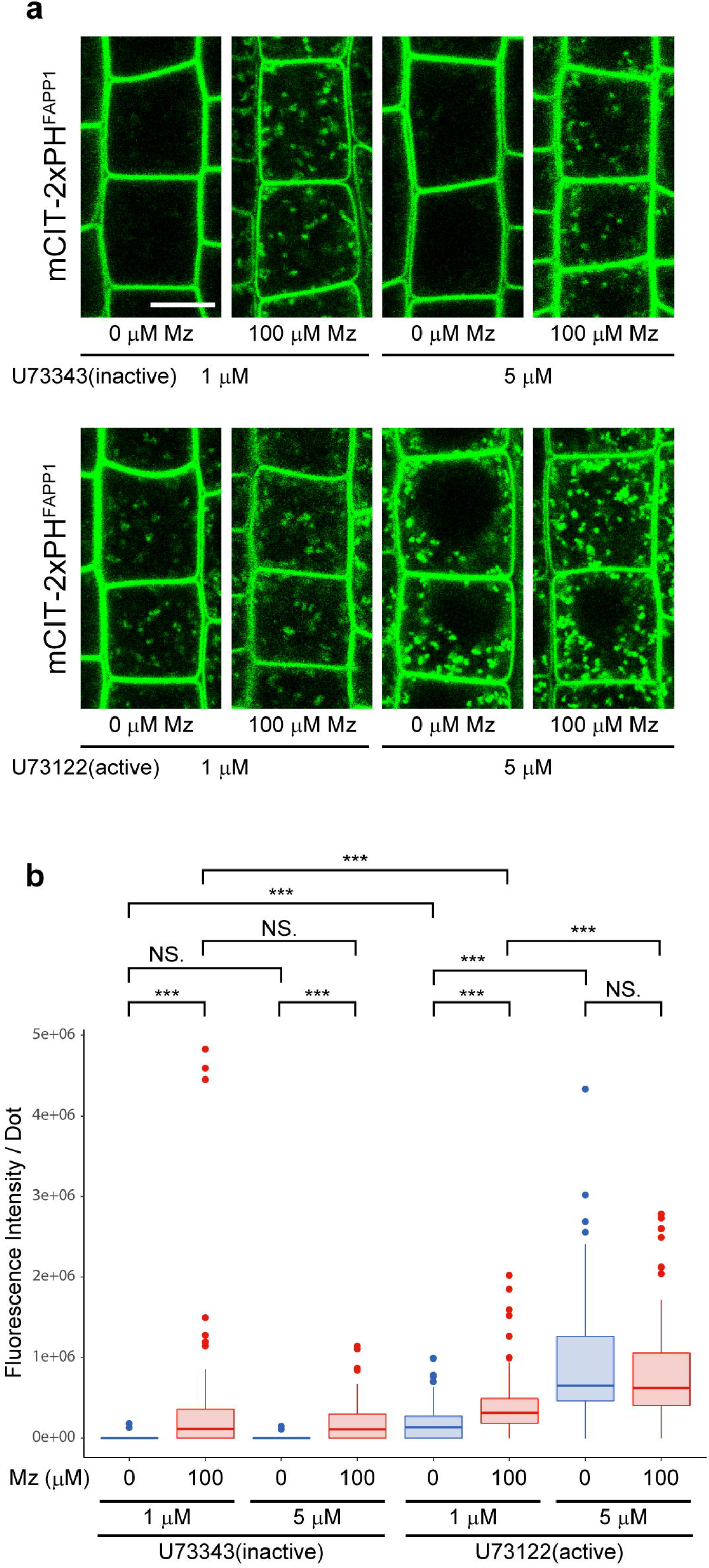
A PI-PLC activity mediates the effect of acyl-chain length of SL on PI4P at the TGN. (a) Confocal images of root epidermal cells expressing the mCIT-2xPH^FAPP1^ PI4P biosensor treated with either the PI-PLC inhibitor U73122 or its inactive analog U73343 at 1 or 5 μM on seedlings grown on 0 or 100 nM Mz-containing medium. (b) Quantification of the fluorescence intensity at intracellular dots (n = more than 60 cells distributed over at least 20 roots for each condition). The PI4P pool at dots was increased upon active U73122 treatment but not with inactive U73343, and this effect was enhanced by increasing the U73122 concentration from 1 to 5 μM. Treatment with 1 μM active U73122 on seedlings grown on 100 nM Mz increased PI4P pool at intracellular dots but not with 5 μM active U73122 indicating that Mz effect is mediated by PI-PLCs. Statistics were done by two-sided Wilcoxson’s ranksum test, *** *P*-value <0.0001. Scale bar, 10 μm.

**Fig. 5.**
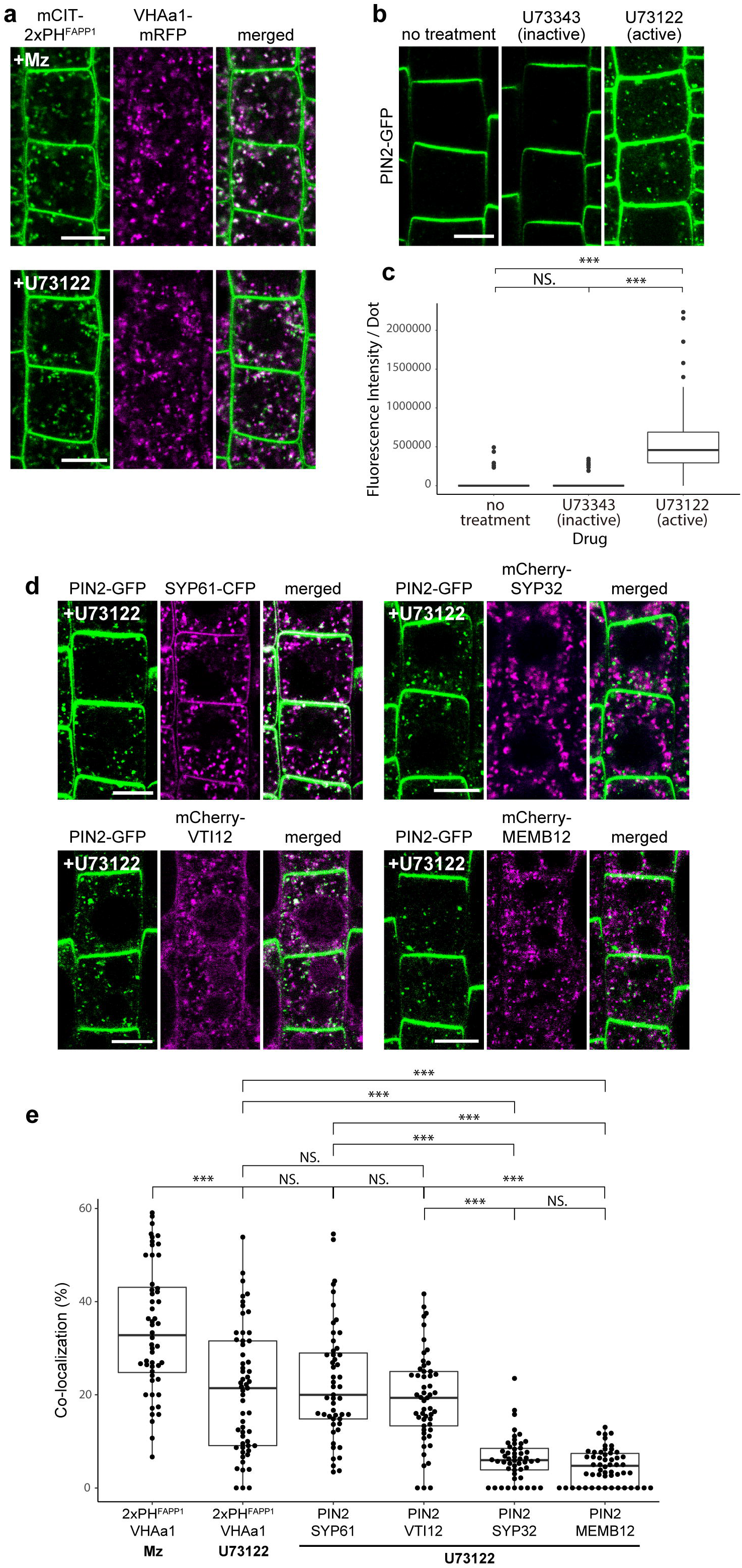
SL-mediated modulation of PI4P through PI-PLC impacts PIN2 sorting at SVs/TGN. (a) Co-localization between endomembrane compartments labeled either by the PI4P biosensor mCIT-2xPHFAPP1 or by the SVs/TGN marker VHA-a1-mRFP. Seedlings were either grown on 100 nM Mz (upper panels) or treated with 5 μM PI-PLC inhibitor U73122 treatment (lower panels). In both cases (Mz or U73122 treatment), the PI4P-positive intracellular dots showed strong co-localization with TGN. (b) Confocal images of root epidermal cells expressing PIN2-GFP upon control condition (left), with 5 μM inactive analog of PI-PLC inhibitor (U73343, middle), and 5 μM active PI-PLC inhibitor (U73122, right). (c) Quantification of the fluorescence intensity of PIN2-GFP specifically at intracellular dots (n = more than 60 cells distributed over at least 20 roots for each condition) showed that PIN2 was accumulated at endomembrane compartments upon U73122 treatment. (d) Co-localization of endomembrane compartments labeled by PIN2-GFP upon 5 μM U73122 treatment with either SVs/TGN labeled by SYP61-CFP (upper left), SVs/TGN labeled by mCherry-VTI12 (lower left), Golgi apparatus labeled by mCherry-SYP32 (upper right), or Golgi apparatus labeled by mCherry-MEMB12 (lower left). (e) Colocalization quantification of (a) and (d) (n = more than 50 cells distributed over at least 20 roots). PIN2 showed significantly higher co-localization with SVs/TGN markers compared to the Golgi markers. Statistics were done by two-sided Wilcoxon’s rank-sum test, *** *P*-value <0.0001. Scale bar, 10 μm.

## Discussion

Homeostasis of PIPs at the TGN is crucial for the regulation of membrane trafficking. In this study, we show that the reduction of the acyl-chain length of SLs results in increased PI4P at the TGN, a main station for sorting of cargos. The interplay between SLs and PIPs has been evidenced in mammals at ER/TGN contact sites and involves a complex regulatory homeostatic loop based on OSBP-mediated PI4P/sterols or PI4P/PS lipid exchange mechanisms induced by the grafting of a polar head on ceramide at the TGN^8,9,15^. We found the whole set of ceramide-based GIPC-synthesizing enzymes in our SVs/TGN proteomics but none of these enzymes were targeted by modification of the acyl-chain length of SL. Since metazachlor does not affect the total amount of SLs^30^, these results strongly support that the acyl-chain length of SLs does not impact the synthesis of SLs. Very long chain FAs (VLCFAs) of 24 or 26 atoms of carbons are predominantly present in the pool of SLs, up to 85% in *Arabidopsis* root, while they constitute less than 2% of the pool of phospholipids and are present mostly in PS^30,62^. Although metazachlor reduces VLCFAs in a large majority in the pool of SLs, it still reduces VLCFAs in the pool of phospholipids as well^30^. There are several lines of evidence indicating that PS is not involved in the interplay between PI4P and SLs at the TGN. First, the localization patterns of both the 2xPH^FAPP1^ and P4M^SiDM^ PI4P biosensors were not drastically altered in the *phosphatidylserine synthase1* (*pss1*) mutant, which does not produce any PS^33^. Second, by contrast to the shortening of the acyl chain length of SLs by Mz treatment, the absence of PS in *pss1* did not induce intracellular PIN2-GFP accumulation or loss of PIN2 polarity^63^. Third, we did not alter the localization of C2^LACT^/PS biosensor upon metazachlor treatment. Fourth, the abundance of PS-related enzymes was not modified in SVs/SYP61 proteomics upon metazachlor. Fifth, we did not observe any change in sterols or PS quantity at the TGN upon metazachlor treatment, which pulls aside an involvement of PIPs/sterols or PIPs/PS exchange mechanism. Hence, the interplay between SLs and PIPs should originate from another mechanism than lipid exchange induced by SL metabolic flux.

Instead, our analyses support a role of PI-PLC in the SL-mediated effect on PI4P homeostasis at the TGN. First, a significant abundance of PI-PLC was detected by proteomics at SVs/SYP61 purified compartments, which is consistent with a previous membrane fractionation report of PLC2 being mostly localized at the PM but as well in the microsomal fraction and small dots inside the cell^53,64^. Second, the abundance of PI-PLC at the TGN was strongly reduced by metazachlor, which fits with the increase of PI4P. Third, the reduction of acyl-chain length of SLs or the inhibition of PI-PLC both led to increased PI4P at the TGN. Fourth, the combined reduction of the acyl-chain length of SLs by metazachlor and the inhibition of PI-PLC did not lead to a simple additive increase of PI4P at the TGN. PI-PLCs are well recognized to catalyze hydrolysis of PI(4,5)P2, but *Arabidopsis* PLCs hydrolyze equally well both PI4P and PI(4,5)P2 *in vitro*, and PI4P is much more abundant *in vivo* compared to PI(4,5)P2^35,54,55^. Moreover, the involvement of PI-PLCs in modulating the global quantity of PI4P has also been previously reported *in planta*^53^. Our results have now uncovered an unexpected role of PLCs in PI4P homeostasis at the TGN. PI-PLC proteins do not have transmembrane domains, but they are known to associate with membrane phospholipids through C-terminal C2 domains and Pleckstrin Homology (PH) domain^52^. However, plant PLCs lack the PH domain which allows the binding of PLC to PIPs in mammals^52,65,66^. Our results reveal that SLs are crucial membrane determinants for modulating the amount of PI-PLC at the plant TGN. We additionally show that modulation of PI4P through PI-PLCs is instrumental in sorting of the auxin efflux carrier PIN2 at SVs of TGN. Consistently with our findings, both acyl chain length of SLs and PI-PLCs are known to be involved in PIN2 polarity, auxin distribution and root gravitropism^30,56,60,61^. Altogether, our work provides a new mechanistic connection between SLs and PI4P at the TGN through SL-mediated consumption of PI4P by PI-PLC. Although the involvement of PI-PLC in protein sorting from the TGN to the PM has been proposed in animal cells^67,68^, their integration in lipid interplay has never been hypothesized while regulation of PI4P quantity at the TGN is instrumental in the face of extensive membrane trafficking and fluctuating lipid metabolic fluxes. Whether this mechanism is common to eukaryotic organisms represents novel perspectives in understanding the function of lipid interplay during protein sorting and cell polarity.

## Methods

### Plant material and growth conditions

The following A. thaliana transgenic fluorescent protein marker lines were used: PI4P biosensors pUB10::mCITRINE-1xPH^FAPP1^ (P5Y, ref^35^), pUB10::mCITRINE-2xPH^FAPP1^ (P21Y, ref^35^), pUB10::mCITRINE-3xPH^FAPP1^ (ref^35^), pUB10::mCITRINE-1xPH^FAPP1^-E50A (ref^32^), pUB10::mCITRINE-1xPH^FAPP1^-E50A-H54A (ref^32^), and pUB10::mCITRINE-P4M^SidM^ (ref^32^). PS biosensor pUB10::mCITRINE-C2^LACT^ (ref^32^). TGN markers pARF1::ARF1-GFP (ref^37^), pSYP61::SYP61-CFP (ref^69^), pVHA-a1::VHA-a1-mRFP (ref^59^), and pUB10::mCherry-VTI12 (W13R, ref^70^). Golgi markers pUB10::mCherry-SYP32 (W22R, ref^70^) and mCherry-MEMB12 (W127R, ref^70^).

Auxin efflux carrier PIN2 marker pPIN2::PIN2-GFP (ref^37^). The double fluorescent lines used for colocalization analysis were established by crossing of the lines above. For confocal observations, sterilized seeds were kept at 4 ^o^C in water for 2-3 days, sown on half Murashige and Skoog (MS) agar medium plates (0.8% plant agar, 1% sucrose, and 2.5 mM morpholinoethanesulfonic acid pH5.8 with KOH), and grown in 16 h light/8 h darkness for 5 days. The growth condition for membrane compartment immunoprecipitation is described hereafter.

### Inhibitor treatments

Metazachlor treatment was performed on seedlings grown for 5 days on half MS plates containing 50 or 100 nM metazachlor (Cayman Chemical). Metazachlor was added to the medium from a 100 mM stock in dimethylsulfoxide by using an intermediate diluted stock at 100 μM (extemporarily prepared).

Wortmannin (Sigma-Aldrich) treatment was performed on seedlings grown on drug-free half MS plates for 5 days and transferred in liquid half MS medium containing 33 μM of wortmannin for the indicated time. PI-PLC inhibitor treatment (U73122 and its inactive analog U73343, Sigma-Aldrich) was performed on seedlings grown on drug-free half MS plates for 5 days and transferred in liquid half MS medium containing 1 or 5 μM of either U73122 or U73343 for 90 min. The drug concentrations are also indicated in the figure legends.

### Confocal microscopy and image analyses

Confocal laser scanning microscopy was performed using a Zeiss LSM 880. Seedlings were mounted with 1/2 MS medium (with or without drugs). Double-sided tape was used as the spacer to separate the slide glass and coverslip. All acquisitions were done with an oil-immersion x40 objective, 1.3 numerical aperture (APO 40x/1.3 Oil DIC UV-IR).

Quantification of the fluorescence intensity was performed by ImageJ. For the PM, the outline of the cell was drawn outside of the PM by hand, and the signal intensity was quantified in the region that is within 1.5 μm inside the outline, which was subsequently normalized by the area. For the intracellular dots, the dots were extracted from the cytoplasmic area by applying a threshold, and their total signal intensity was normalized by the number of the dots. The threshold was kept constant for all the samples that are shown in the same graph. In order to avoid including non-dotty background, only the structures with the circularity over 0.1 were quantified (circularity is defined as *4πA/P^2^* with *A*=area and *P*=perimeter, and it takes the values 0.0-1.0 with 1.0 represents the perfect circle). For the experiments using PI-PLC inhibitor (Fig. 4b and 5c, e), a size filter of 10-400 pixels (approximately 0.1-4 μm^2^) was additionally applied.

Co-localization analyses were performed using the geometrical (centroid) object-based method^71^. The cells were segmented by hand and subcellular compartments were extracted by applying a threshold, and the distance between the centroids of two objects was calculated using 3D objects counter plugin of imageJ. The threshold was kept constant between samples of the same fluorescent line. When the distance between two labelled structures was below the optical resolution limit, the co-localization was considered as true. The resolution limit was calculated based on the shorter emission maximum wavelength of the fluorophores. For the better extraction of the subcellular compartments, a size filter of 10-400 pixels (approximately 0.1-4 μm^2^) was applied.

### Statistical analyses

All statistical analyses are two-sided Wilcoxon’s rank-sum test, and they were performed with R (version 3.6.0) and RStudio (version 1.2.1335). Variances between each group of data are represented in boxplot, bee swarm plot, or s.d.. *P*-values and sample sizes are described in the figure legends.

### Immunoprecipitation of intact SVs/TGN

Immunoprecipitation of intact SVs/TGN membrane compartments was performed as described^30,72^. In brief, Arabidopsis seedlings were grown in flasks with liquid half MS medium for 9 days under 120 rpm, shaking and 16 h light/8 h darkness cycle. Seedlings were grinded by an ice-cooled mortar and pestle in vesicle isolation buffer (HEPES 50 mM pH 7.5, 0.45 M sucrose, 5 mM MgCl2, 1 mM dithiothreitol, 0.5% polyvinylpyrrolidone, and 1 mM phenylmethylsulfonyl fluoride). The homogenate was filtered and centrifuged to remove the debris. The supernatant was loaded on 38% sucrose cushion (dissolved in 50 mM HEPES pH 7.5) and centrifuged at 150,000 *g* for 3 h at 4 °C. After removing the supernatant above the membrane pool located at the interface between the sucrose and the supernatant, a step-gradient sucrose was built on the top of the membrane interface with successive 33% and 8% sucrose solutions (dissolved in 50 mM HEPES pH 7.5). The tubes were centrifuged overnight at 150,000 *g* at 4 °C, the membranes that appeared at the 38/33% and 33/8% sucrose interfaces were harvested, pooled and diluted in 2-3 volume of 50 mM HEPES pH 7.5. After a centrifugation step at 150,000 g for 3 h at 4 °C, the pellet was resuspended in resuspension buffer (50 mM HEPES pH 7.5, 0.25 M sucrose, 1.5 mM MgCl_2_, 150 mM NaCl, 1 mM phenylmethylsulfonyl fluoride, and protease inhibitor cocktail from Sigma-Aldrich). The protein amount of the resuspended membrane fractions was quantified by Bicinchoninic Acid Protein Assay Kit (Sigma-Aldrich) and equilibrated between different samples. Those equilibrated membrane fractions were used as the IP input. IP was performed with magnetic Dynabeads coupled to proteinA (Thermo Fisher Scientific) conjugated with rabbit anti-GFP antibody (Thermo Fisher Scientific, A-11122) by bis[sulfosccinimidyl] suberate (Thermo Fisher Scientific, 21580). The beads were incubated with the IP input for 1 h at 4 °C, washed and resuspended in resuspension buffer.

### Western blotting of IP fractions

To equally load the IP input and the beads-IP fraction (IP output) for Western blotting, TGX Stain-Free FastCast premixed acrylamide solution (Bio-Rad) and ChemiDoc MP imaging system (Bio-Rad) were used to visualize the proteins. The whole individual lanes were quantified by ImageJ software and the quantity of proteins were adjusted to get equal new loading between each lane. Western blotting was performed with mouse anti-GFP (1/1,000, Roche, 118144600001) and rabbit anti-ECH^73^ (1/1,000) as the primary antibodies, and goat anti-mouse IgG-HRP conjugate (1/3,000, Bio-Rad, 1721011) and anti-rabbit IgG-HRP conjugate (1/5,000, Bio-Rad, 1706515) as the secondary antibodies.

### Characterization of lipid composition

Lipid characterization of immuno-precipitated intact TGN compartments is fully described in^30,72^. Shortly, for the characterization of fatty acids, 50 μl of IP beads fraction was incubated with 1 ml of 5% sulfuric acid in methanol, including 5 μg/ml of the lipid standards C17:0 to normalize non-hydroxylated fatty acids and h14:0 to normalize α-hydroxylated fatty acids. The transesterification was performed overnight at 85°C. After cooling down at room temperature, the fatty acids methyl esters (FAMEs) were extracted by adding 1 ml of NaCl 2.5% and 1 ml of hexane. After hand shaking and centrifugation at 700 *g* for 5 min at room temperature, the higher phase was collected and placed in a new tube. After addition of 1 ml of 100 mM Tris, 0.09% NaCl pH 8 with HCl, hand shaking and centrifugation, the higher phase was collected and evaporated. After evaporation, 200 μl of *N*,*O*-Bis(trimethylsilyl)trifluoroacetamide+1% trimethylsilyl (BSTFA+1% TMCS, Sigma) were added and incubated at 110 °C for 15 min. After evaporation, lipids were resuspended in 80 μl of 99% hexane. For the characterization of sterols, 50 μl of IP beads fraction was incubated with 1 ml of chloroform:methanol 2:1, including 5 μg/ml of the sterol standard α-cholestanol, for 2 h at room temperature. After addition of 1 ml of 0.9% (w/v) NaCl, hand shaking and centrifugation, the lower organic phase was collected and evaporated. Fatty acids were removed by saponification by adding 1 ml of 99% ethanol and 100 μl of 11N KOH for 1h at 80°C. Then, 1 ml of 99% hexane and 2 ml of water were added. After hand shaking and centrifugation, the higher phase was collected, placed in a new tube where 1 ml of 100 mM Tris, 0.09% NaCl pH 8 with HCl. After hand shaking and centrifugation, the higher phase was collected, placed in a new tube and evaporated.

After evaporation, 200 μl of BSTFA+1% TMCS were added and incubated at 110 °C for 15 min. After evaporation, lipids were resuspended in 80 μl of 99% hexane. GC-MS was performed using an Agilent 7890A and MSD 5975 Agilent EI with the following settings: the helium carrier gas was set at 2 ml/min, the splitless mode was used for injection, the temperatures of injector and auxiliary detector were set at 250°C and 352°C respectively, the oven temperature was held at 50°C for 1 min, a 25°C/min ramp (2-min hold) and a 10°C/min ramp (6-min hold) were programmed at 150°C and 320°C respectively, the MS analyzer was set in scan only with a mass range of 40-700m/z in positive mode with electron emission set to 70 eV, the MS source and the MS Quad were set to 230°C and 50°C respectively.

### Label-free LC-MS/MS proteomic analysis of SVs/TGN compartments

Proteins of the IP output fraction were eluted by adding 25 μL of 1% SDS, 0.3 μL of 2M dithiothreitol, 2.3 μL of 1M iodoacetamide, and 6.9 μL of 5x Laemmli buffer sequentially (the volume of each reagent is for 75 μL Dynabeads in initial amount) with an incubation for 30 min at 37 °C (except for iodoacetamide at the room temperature) between each addition. The protein amounts of the eluted samples were equilibrated using the Stain-Free protein visualization system similarly to the loading controls for Western blotting described above. Samples from four biological replicates were used for quantification. The equilibrated samples were solubilized in Laemmli buffer and deposited onto SDS-PAGE gel for concentration and cleaning purposes. After colloidal blue staining, bands were cut out from the gel and subsequently cut into 1 mm^3^ pieces. Gel pieces were destained in 25 mM ammonium bicarbonate and 50% acetonitrile (ACN), rinsed twice in ultrapure water, and shrunk in ACN for 10 min. After ACN removal, the gel pieces were dried at room temperature, covered with trypsin solution (10 ng/μL in 50 mM NH4HCO3), rehydrated at 4 °C for 10 min, and finally incubated overnight at 37 °C. Gel pieces were then incubated for 15 min in 50 mM NH4HCO3 at room temperature with rotary shaking. The supernatant was collected, and an H2O/CAN/HCOOH (47.5:47.5:5) extraction solution was added onto gel slices for 15 min. The extraction step was repeated twice. Supernatants were pooled and concentrated in a vacuum centrifuge to a final volume of 100 μL. Digests were finally acidified by addition of 2.4 μL of formic acid (5% v/v).

Peptide mixture was analyzed on an Ultimate 3000 nanoLC system (Dionex) coupled to an Electrospray Q-Exactive quadrupole Orbitrap benchtop mass spectrometer (Thermo Fisher Scientific). 10 μL of peptide digests were loaded onto a C18 PepMap trap column (300 μm inner diameter x 5 mm, Thermo Fisher Scientific) at a flow rate of 30 μL/min. The peptides were eluted from the trap column onto an analytical C18 PepMap column (75 μm inner diameter x 25 cm, Thermo Fisher Scientific) with a 4-40% linear gradient of solvent B in 108 min (solvent A was 0.1% formic acid in 5% ACN, and solvent B was 0.1% formic acid in 80% ACN). The separation flow rate was set at 300 nL/min. The mass spectrometer was operated in positive ion mode at a 1.8 kV needle voltage. Data were acquired using Xcalibur 2.2 software in a data-dependent mode. MS scans (m/z 350-1600) were recorded at a resolution of R=70,000 (m/z 200) and an AGC target of 3×10^6^ ions collected within 100 ms. Dynamic exclusion was set to 30 s and top 12 ions were selected from fragmentation in HCD mod. MS/MS scans with a target value of 1×10^5^ ions were collected with a maximum fill time of 100 ms and a resolution of R=175,000. Additionally, only +2 and +3 charged ions were selected for fragmentation. The other settings were as follows: no sheath nor auxiliary gas flow, heated capillary temperature at 250 °C, normalized HCD collision energy of 25%, and an isolation width of 2 m/z.

Data were searched by SEQUEST through Proteome Discover 1.4 (Thermo Fisher Scientific) against Araport v11 protein database. Spectra from peptides higher than 5000 Da or lower than 350 Da were rejected. The search parameters were as follows: mass accuracy of the monoisotopic peptide precursor and peptide fragments was set to 10 ppm and 0.02 Da respectively. Only b- and y-ions were considered for mass calculation. Oxidation of methionines (+16 Da) was considered as variable modification and carbamidomethylation of cysteines (+57 Da) as fixed modification. Two missed trypsin cleavages were allowed. Peptide validation was performed using Percolator algorithm^74^ and only “high confidence” peptides were retained corresponding to a 1% False Positive Rate at peptide level.

For label-free quantitative data analysis, raw LC-MS/MS data were imported in Prigenesis QI for Proteomics 2.0 (Nonlinear Dynamics). Data processing includes the following steps: (i) features detection. (ii) features alignment across the samples to compare, (iii) volume integration for 2-6 chargestate ions, (iv) normalization on features ratio median, (v) import of sequence information, (vi) calculation of protein abundance (sum of the volume of corresponding peptides), (vii) a t-test to compare each group and filtering of proteins based on *p*-value <0.05. Only non-conflicting features and unique peptides were considered for calculation at protein level.

## Acknowledgments

We are grateful to Sebastien Mongrand (LBM, Bordeaux, France) and Rishi Bhalerao (UPSC, Umeå, Sweden) for critical reading of the manuscript. We thank Fabrice Cordelières (Bordeaux Neurocampus, France) for discussions and advices on microscopy images quantification and Clément-Marie Train (Swiss Institute of Bioinformatic, Lausanne, Switzerland) for his help on proteomic data. Lipidomic analyses were performed on the Bordeaux Metabolome Facility-MetaboHUB (ANR-11-INBS-0010). Imaging was performed on the plant unit of the Bordeaux Imaging Center (BIC). This work was supported by a PhD funding from the French ministry of research (MENRT) granted to Y.B. to support the PhD thesis of N.E, Overseas Research Fellowship granted from Japan Society for Promotion of Science (JSPP) to Y.I., a research grant from the French National Research Agency (ANR) to Y.B and Y.J (ANR-18-CE13-0025), and a research grant from the European Research Council (ERC) to Y.J. (336360-APPL).

## Author contributions

Y.B. conceptualized and designed the experiments with input from Y.J. and P.M. Y.I. performed experiments in Fig. 1, 2, 4, 5. N.E. generated material for proteomic experiment in Fig. 3, S.C. performed LC-MS proteomics of Fig. 3, Y.I, N.E and Y.B. analyzed proteomic results. Y.B. and W. M. performed lipid extraction and analysis in Fig. 2. M.P. and L.N. provided crossed lines useful for this study. All quantifications and statistics were performed by Y.I. Y.I. built all the figures. Y.B. wrote the manuscript with input from all the authors. Y.B. acquired funding and supervised all aspect of the study. All authors reviewed, edited and approved the manuscript.

---

**Supplementary Data 1. Full list of proteins found in the LC-MS/MS label-free quantitative proteomics of SVs/TGN immuno-purified compartments.** This is the complete dataset for the proteomic analysis of SYP61-SVs/TGN compartments presented in Fig. 3 and Supplementary Fig. 3. Abundance values were calculated for each accession found (the number of peptides found is indicated) in each of the four biological repeats for both control condition and metazachlor (Mz) treatment. Average abundances for both control and Mz treatment and the ratio between Mz treatment and control were calculated resulting in a deprivation or enrichment value of a given protein in SYP61-SVs/TGN compartment upon metazachlor treatment.

**Supplementary Data 2. Short list of proteins displayed in this study and found in proteomics of SVs/TGN immuno-purified compartments.** This is an extracted dataset of the proteins displayed in Fig. 3 and Supplementary Fig. 3 from Supplementary Data 1. Abundance values, average abundances for both control and Mz treatment and the ratio between Mz treatment and control are presented as in Supplementary Data 1.

**Supplementary Fig. 1. The acyl-chain length of SL is required for the intracellular distribution of PI4P independently of ARF1.** (a) Confocal micrographs of root epidermal cells expressing the PI4P sensor mCIT-1xPH^FAPP1-E50A-E54A^ upon Mz. Fluorescence intensity at intracellular dots (b) and at the PM (c) was quantified (n = more than 60 cells distributed over at least 20 roots for each experiment). (d) Confocal micrographs of root epidermal cells expressing GFP-tagged ARF1. Fluorescence intensity at intracellular dots was quantified (e, n =more than 20 roots for each experiment). (f) Confocal micrographs of root epidermal cells expressing the PI4P sensor mCIT-P4M^SidM^ upon Mz. The fluorescence intensity at the PM was quantified (g, n = more than 60 cells distributed over at least 20 roots for each experiment). Statistics were done by two-sided Wilcoxson’s rank-sum test, *** *P*-value <0.0001. Scale bars, 10 μm.

**Supplementary Fig. 2. Detailed GC-MS analysis of fatty acids and sterols at immuno-purified SVs/TGN fraction with or without Mz treatment.** This is the detailed analysis of Fig. 2d. (a) Mz treatment strongly reduces SL-specific α-hydroxylated fatty acids h24:1, h24:0 and h26:0 and non-α-hydroxylated fatty acids C24:1, C24:0 and C26:0 (present in both sphingolipids and PS). (b, c) The pool of C16 and C18, mainly present in phospholipids, is not altered by Mz as shown in details (b) or by the sum of C16 or the sum of C18 (c). n = 3 biological replicates, error bars are s.d..

**Supplementary Fig. 3. Immuno-purified SVs/TGN fraction is enriched for SVs/TGN markers SYP61 and ECH and contain a set of SL- and PS-related enzymes not targeted by acyl-chain length of SLs.** (a-c) Immuno-purification of SYP61 SVs/TGN compartments for label-free proteomics. (a) Loading control for (b) and (c). IP input and output were loaded on Stain-Free polyacrylamide gel to confirm their equal loading. (b, c) Western-blotting images with anti-GFP (b) or anti-ECHIDNA (c) antibodies showing enrichment of SVs/TGN compartments in IP output over the microsomal IP input. (d-f) Proteomics results of sphingolipid or PS related enzymes. (d) Protein abundance of sphingolipid related enzymes. (e) Protein abundance of PS related enzymes. (f) The abundance ratio of the proteins shown in (d) and (e) between control and Mz treated sample. The same threshold as Figure 3 was applied.

