## Supplementary figures and images for "Sphingolipids mediate polar sorting of PIN2 through phosphoinositide consumption at the *trans*-Golgi Network"

### Supplementary Figure 1

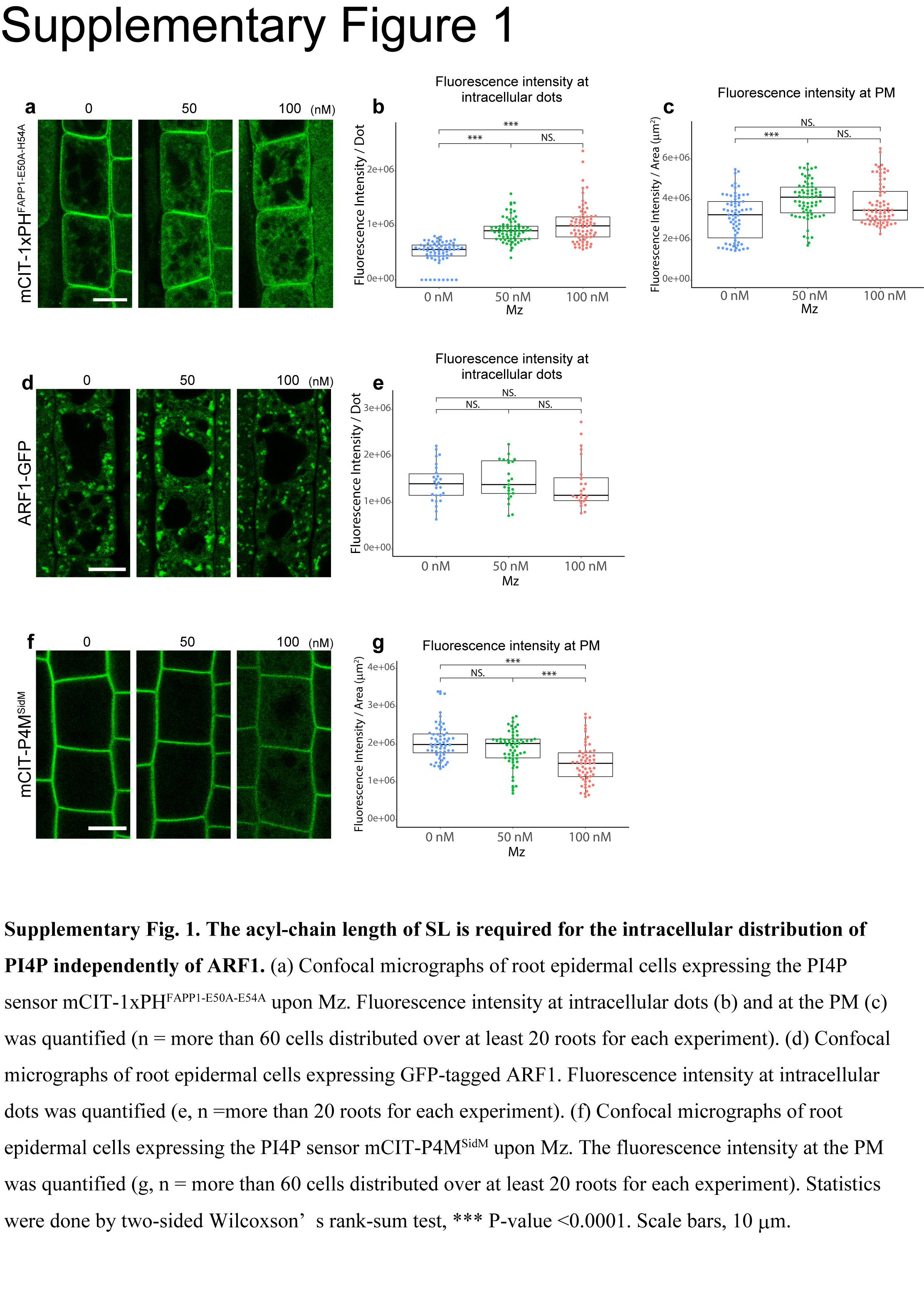

### Supplementary Figure 2

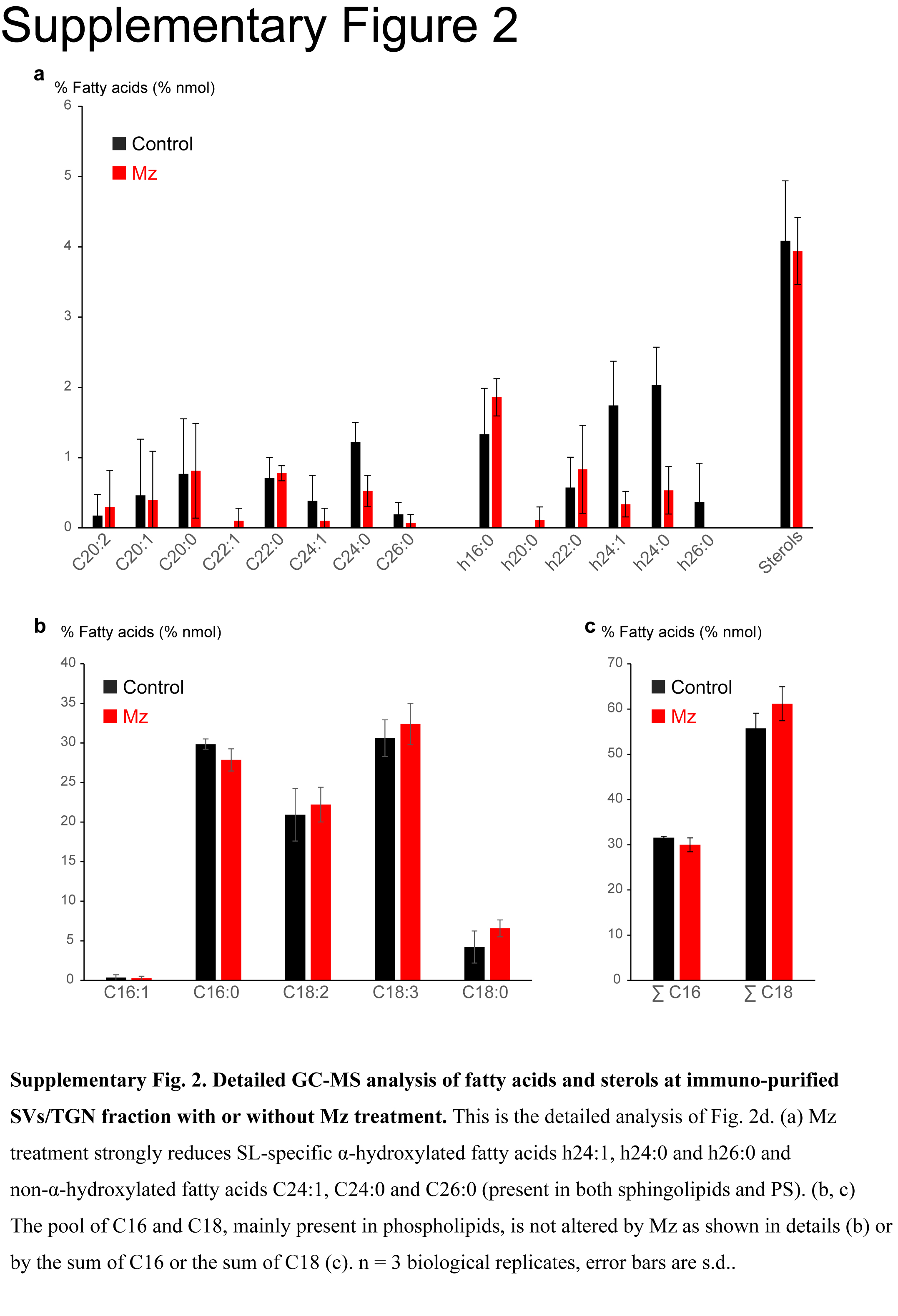

### Supplementary Figure 3

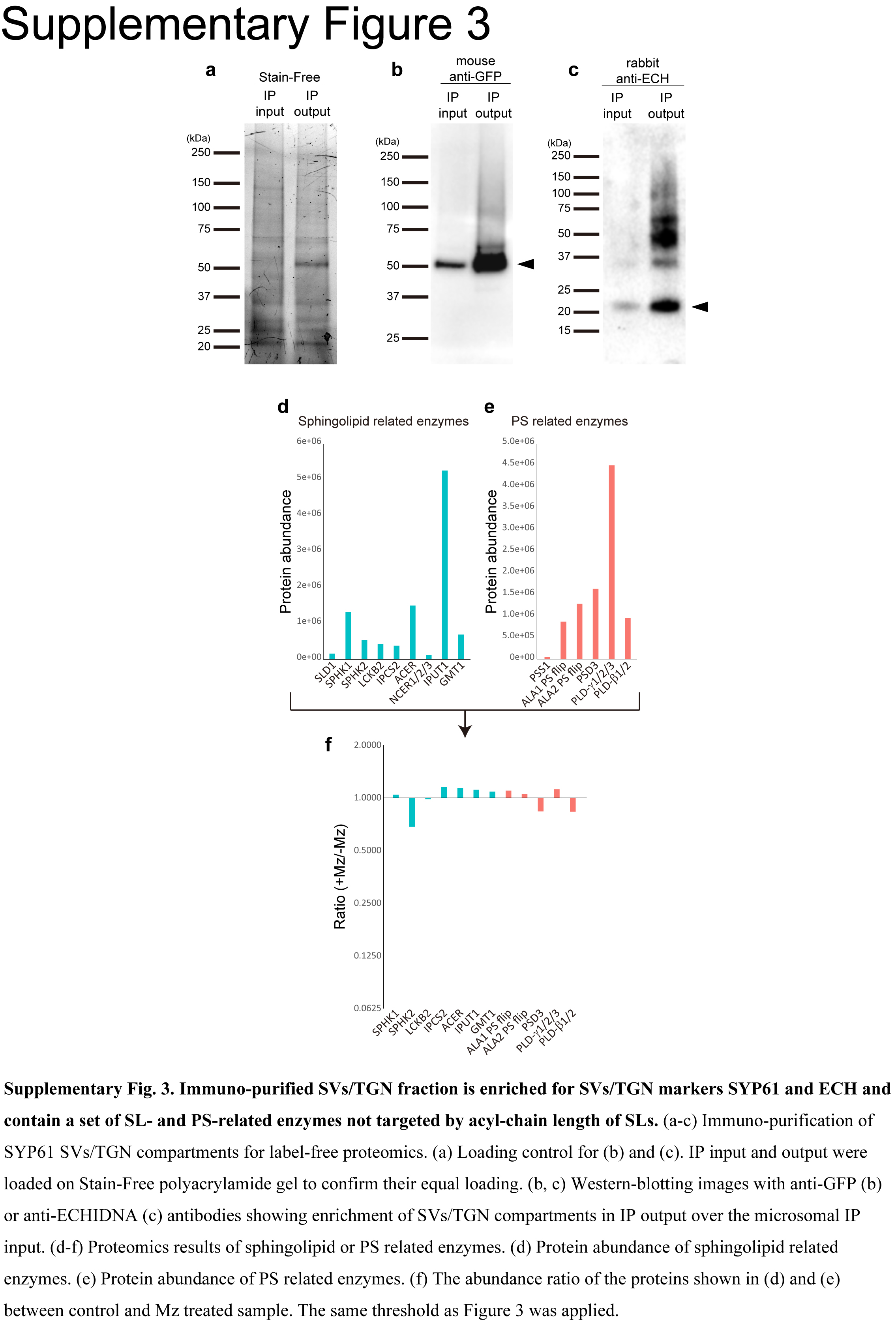
